# Capturing actively produced microbial volatile organic compounds from human associated samples with vacuum assisted sorbent extraction

**DOI:** 10.1101/2021.02.16.431476

**Authors:** Joann Phan, Joseph Kapcia, Cynthia I. Rodriguez, Victoria L. Vogel, Sage J. B. Dunham, Katrine Whiteson

## Abstract

Volatile organic compounds (VOC) from biological samples have unknown origins. VOCs may originate from the host or different microbial organisms from a microbial community. In order to disentangle the origin of microbial VOCs, we performed volatile headspace analysis of bacterial mono- and co-cultures of *Staphylococcus aureus, Pseudomonas aeruginosa*, and *Acinetobacter baumannii*, and stable isotope probing in biological samples of feces, saliva, sewage, and sputum. We utilized mono- and co-cultures to identify volatile production from individual bacterial species or in combination with stable isotope probing to identify the active metabolism of microbes from the biological samples. To extract the VOCs, we employed vacuum assisted sorbent extraction (VASE). VASE is an easy-to-use commercialized solvent-free headspace extraction method for semi-volatile and volatile compounds. The lack of solvents and the near vacuum conditions used make developing a method relatively easy and fast when compared to other extraction options like tert-butylation and solid phase microextraction. However, VASE does not work on nonvolatile compounds, thus excluding many protein analyses and heavy stable isotope labelling experiments. Using the workflow described here, we identified volatile signatures from mono- and co-cultures indicating there were volatiles specific to certain microbes or co-cultures. Furthermore, analysis of the stable isotope probing of biological samples identified VOCs that were either commonly or uniquely produced from the different human derived biological samples. Here we present the general workflow and experimental considerations of VASE.

**SUMMARY:** From this protocol, readers will be able to extract volatile organic compounds from a biological sample with the vacuum assisted sorbent extraction method, run samples on a GC-MS with the use of the Entech autosampler, and analyze data.

## INTRODUCTION

Volatile organic compounds (VOCs) have great promise for bacterial detection and identification because they are emitted from all organisms and different microbes have unique VOC signatures. Volatile molecules have been utilized as a non-invasive measurement for detecting various respiratory infections including chronic obstructive pulmonary disease ^1^, tuberculosis^2^ in urine^3^, and ventilator associated pneumonia ^4^, in addition to distinguishing subjects with cystic fibrosis (CF) from healthy control subjects^5,6^. Volatile signatures have even been able to distinguish specific pathogen infections in CF (*Staphylococcus aureus* ^7^, *Pseudomonas aeruginosa*, ^8,9^, and *S. aureus vs. P. aeruginosa*, ^10^). With the complexity of such biological samples, it is difficult to pinpoint the source of the important volatile molecules.

One strategy for disentangling the volatile profiles from multiple infecting microbes is performing headspace analysis of strains in both mono- and co-culture ^11^. Headspace analysis examines the analytes emitted from a sample rather than within or embedded in the sample itself. Microbial metabolites have often been characterized in mono-cultures because it is much more difficult to determine the origin of microbial metabolites in complex clinical samples. By profiling volatiles from bacterial mono-cultures, the types of volatiles a microbe produces in vitro may represent a baseline of its volatile repertoire. Combining bacterial cultures, e.g., creating co-cultures, and profiling the volatile molecules produced by co-cultures may reveal the interactions or cross-feeding between the bacteria^12^.

Another strategy for identifying the microbial origin of volatile molecules is to provide a nutrient source that is labeled with a stable isotope. Stable isotopes are naturally occurring non-radioactive forms of atoms with a different number of neutrons. The utilization of stable isotopes has been used since the early 1930s to trace active metabolism in animals ^13^. More recently, a stable isotope in the form of heavy water (D_2_O) has been used to identify metabolically active S. aureus in a clinical CF sputum sample ^14^. In another example, ^13^C labeled glucose has been used to demonstrate the cross-feeding of metabolites between CF clinical isolates of *P. aeruginosa* and *Rothia mucilaginosa*^12^.

With the advancement of mass spectrometry techniques, methods of detecting volatile cues have moved from qualitative observations to more quantitative measurements. By using gas chromatography mass spectrometry (GC-MS), processing of biological samples has become within reach for most laboratory or clinical settings. Many methods to survey volatile molecules have been used to profile samples such as food, bacterial cultures and other biological samples, and air and water to detect contamination. However, several common methods of volatile sampling with high throughput do not involve solvent-less and vacuum extractions.

Here, we employed a method called vacuum assisted sorbent extraction (VASE) followed by thermal desorption on a GC-MS to survey the volatile profiles of bacterial mono- and co-cultures and to identify actively produced volatiles with stable isotope probing from human feces, saliva, sewage and sputum samples (Figure 1). Isotope probing experiments with human samples required adding a stable isotope source, such as ^13^C glucose, and media to cultivate the growth of the microbial community. The active production of volatiles was identified as a heavier isotope molecule by GC-MS. Extraction of volatile molecules under a static vacuum enabled the detection of volatile molecules with increased sensitivity^15–17^.

**Figure 1.**
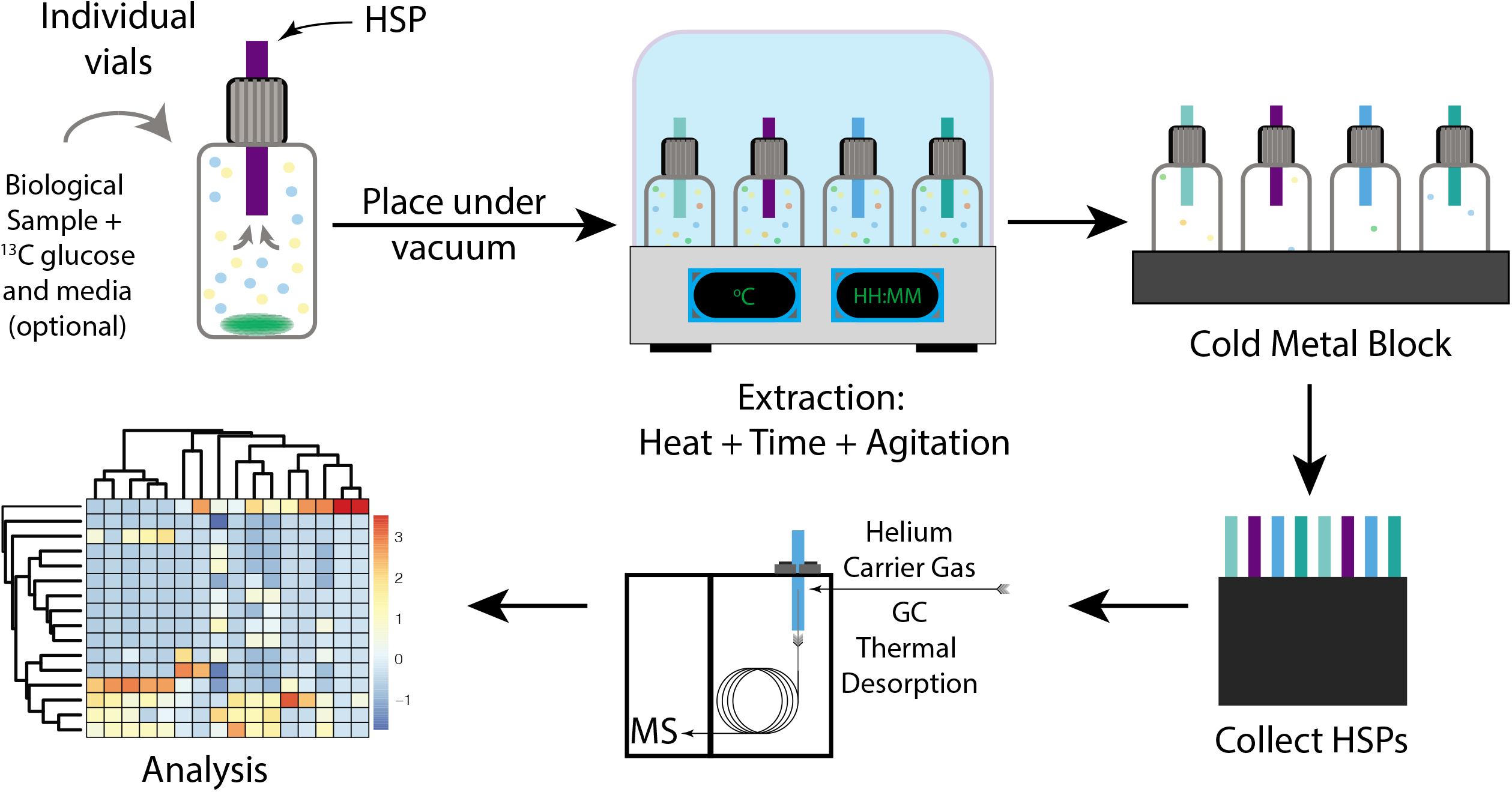
Protocol schematic. A biological sample is placed into a glass vial and assembled with the lid liner and headspace sorbent pen (HSP). A vacuum is applied to the vial until 30 mmHg is reached. The vacuum source is removed, and the vials are placed in the sorbent pen extraction unit (SPEU) where there is a static extraction with heat, agitation with rpm, and time. After extraction, vials are placed on a cold metal block to remove water from the headspace. The HSPs are collected and run via thermal desorption on the GC-MS. The data are then analyzed with ChemStation, DExSI, and R.

## REPRESENTATIVE RESULTS

### Mono- and co-cultures of *S. aureus, P. aeruginosa, and A. baumannii*

The mono- and co-cultures consisted of the bacterial species *S. aureus, P. aeruginosa,* and *Acinetobacter baumannii*. These are common opportunistic pathogens found in human wounds and chronic infections. To identify the volatile molecules present in the mono- and co-cultures, we performed a short 1-hour extraction at 70 °C with 200 rpm agitation. From the mono- and co-cultures at 24- and 48-hour timepoints, we detected 43 annotated volatile molecules (Figure 2). The types of volatile molecules detected were aldehydes, ketones, alcohols, sulfuric compounds, hydrocarbons, carboxylic acids or esters, and aromatics. There were a small number of volatile molecules that were only detected in certain mono- or co-cultures at certain timepoints. For example, volatiles acetoin and 3-hydroxy-2-butanone acetate were only detected in the *S. aureus* cultures at the 48-hour timepoint (Figure 2). Volatile 1-propanol 2-methyl was detected only in the *P. aeruginosa* and *A. baumannii* co-culture at 48 hours (Figure 2). Ethyl acetate was present in *A. baumannii* co-cultures with either *S. aureus* or *P. aeruginosa* at 48 hours (Figure 2). The metabolites heptane, 2,3-dimethyl and pentane, 2-methyl were only detected in the *A. baumannii* culture at 24 hours (Figure 2). Acetaldehyde and ethanol had higher relative abundances in the *A. baumannii* and *S. aureus* co-culture at the 24-hour timepoint compared to 48 hours and either of the strains in culture alone (Figure 2). Some of the volatiles were more abundant in cultures at either the 24- or 48-hour timepoint. Short chain fatty acids, acetic acid, butanoic acid, and propanoic acid were at high relative abundances in cultures at 48 hours but were not detected in the 24-hour cultures (Figure 2). Hexane was more abundant in the TH control at 24 hours compared to 48 hours (Figure 2).

**Figure 2.**
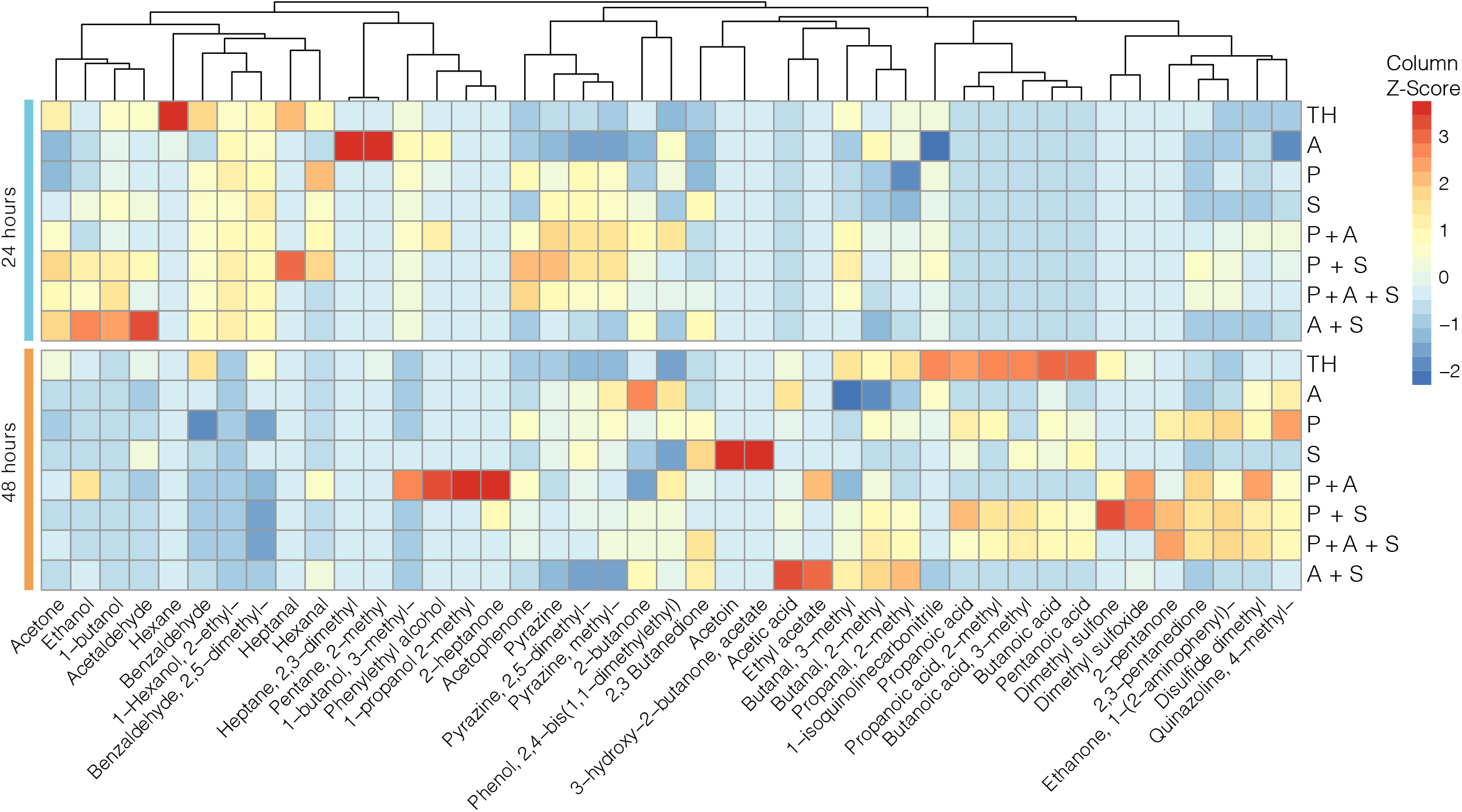
Heatmap of mono- and co-cultures. VOCs detected from mono- and co-cultures at 24- and 48-hour timepoints. The bacterial species are A = *A. baumanii*, P = *P. aeruginosa*, and S = *S. aureus*. The co-cultures are the combinations of the letters representing each strain. The media control is TH = Todd Hewitt media. All samples were extracted for 1 hour at 70 °C with 200 rpm agitation. Heatmap intensity values are column Z-scores, normalized by metabolite. The Z-score was calculated by taking the difference of value and the mean of the value, dividing by the standard deviation of the value. The dendrogram was generated with the “cluster_cols” option in the “pheatmap” function R. The dendrogram represents hierarchical clustering in which metabolites that cluster together have more similar z-score values across samples.

### Stable isotope labeling of fecal, sewage, and saliva samples

To identify active production of volatile molecules from a biological sample, we added a labeled nutrient source, ^13^C glucose or D_2_O, and media to support the growth of the microbial community. We analyzed one unique sample from each of the different sample types of fecal, sewage, and saliva samples in triplicate. There was more incorporation of the ^13^C into fully labeled volatile molecules (Figure 3A-D) compared to incorporation with deuterium (Figure 3E). The ^13^C was incorporated into 2-butanone, 3-hydroxy; 2,3-butanedione; acetic acid; and phenol for all fecal, sewage, and saliva samples (Figure 3A). The other labeled volatiles were detected in either two or one sample types. For example, acetone, butanoic acid, and propanoic acid were detected as labeled in saliva and sewage (Figure 3B). The labeled volatiles dimethyl trisulfide and disulfide dimethyl were enriched in both fecal and saliva samples (Figure 3C). Volatiles 1-propanol, 2-butanone, benzophenone, ethanol, and methyl thiolacetate were enriched only in sewage (Figure 3D). The labeled volatile 2,3-pentanedoine was enriched in saliva (Figure 3D). Deuterium was incorporated into the volatiles acetic acid; benzaldehyde, 4-methyl; dimethyl trisulfide; and phenol from either saliva or sewage samples (Figure 3E). In addition to the isotope enriched volatiles, there were volatiles detected that did not contain incorporated stable isotopes. For example, pyrazine compounds, except for pyrazine, 2,5-dimethyl, were detected in fecal, sewage, and saliva samples but were not fully enriched with ^13^C (Figure S1).

**Figure 3.**
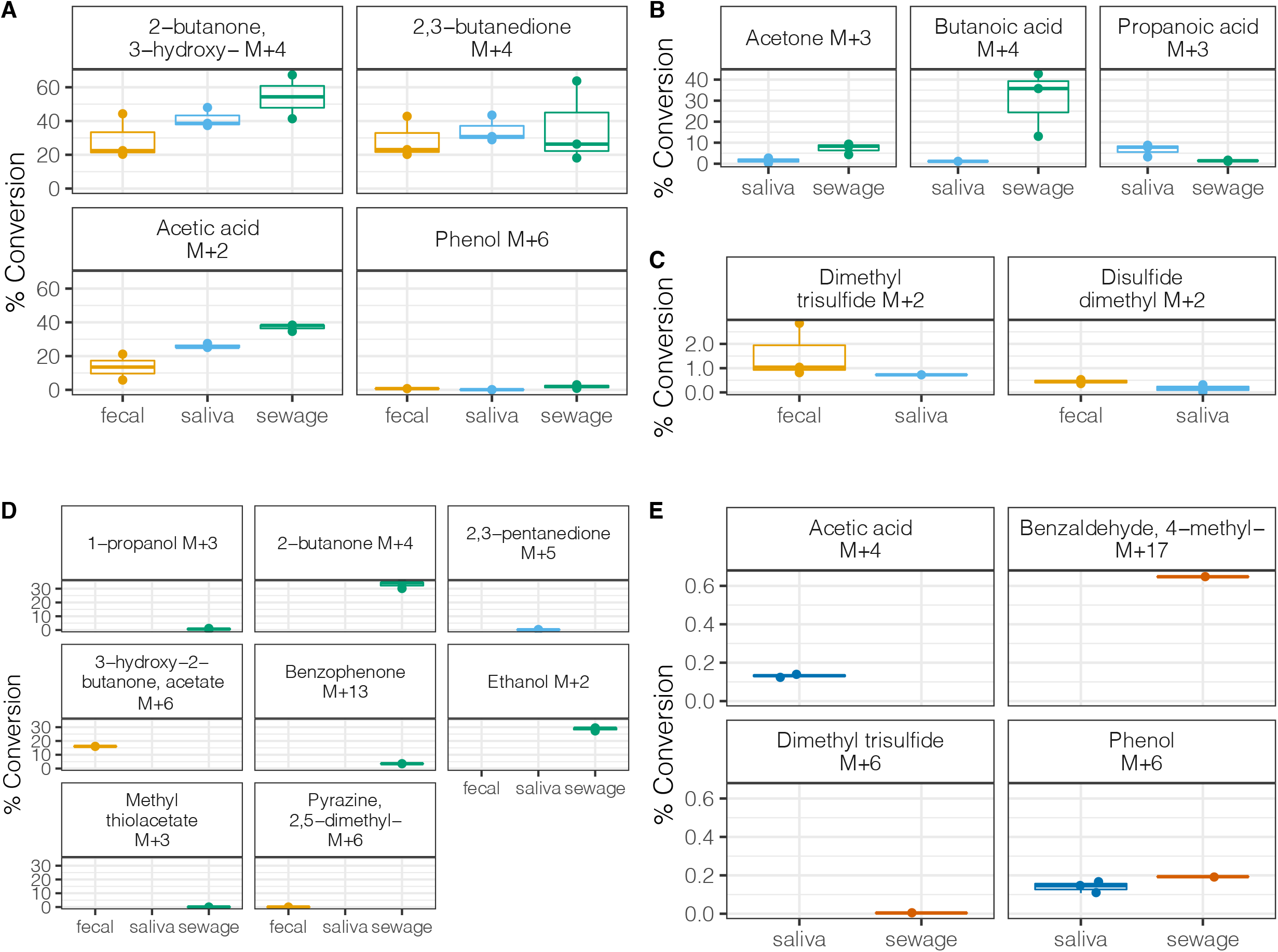
Percent conversion of ^13^C into volatile molecule mass in fecal, saliva, and sewage samples during 18 hours of incubation during extraction. The % conversion was calculated for fully labeled compounds by taking the mass of the fully labeled compound (M+N) and dividing it by (M+N) + the mass of the unlabeled volatile mass (M). N is the maximum number of possible carbons (in A-D) or hydrogens (in E) that can be labeled in each volatile molecule. Fully labeled compounds are when all carbons of the volatile are replaced by ^13^C. Where data are missing, the volatile was not detected. For example in panel (D), 1-propanol was not detected in fecal or saliva samples. Number of replicates per sample = 3. (A) The ^13^C labeled volatiles detected in all sample types (feces, saliva, and sewage). (B) The ^13^C labeled volatiles only detected in saliva and sewage samples. (C) The ^13^C labeled volatiles detected in feces and saliva samples. (D) The ^13^C labeled volatiles detected in one of the three different sample types. (E) The deuterium labeled volatile molecules.

### Stable isotope labeling of sputum samples

We implemented the strategy for identifying actively produced volatiles with sputum samples from seven human subjects with cystic fibrosis. We compared the volatiles in the sample with those that emerged from samples cultured with a stable isotope label. Each sample was analyzed twice: we first extracted volatiles from sputum samples and then performed stable isotope probing with ^13^C glucose and media. The samples collected from the subjects spanned three different timepoints or clinical states, baseline, exacerbation, and treatment^18^. The volatiles detected as labeled in the cultured sputum samples had different relative abundances compared to the unlabeled volatiles from the uncultured sputum samples. Culturing conditions in the stable isotope probing experiments with sputum may favor the growth of certain microbes, leading to differences in relative abundances of volatiles compared to the uncultured sputum samples. For example, acetic acid, dimethyl trisulfide, acetone, and propanal, 2-methyl were more abundant in the cultured sputum samples compared to the uncultured sputum samples (Figure 4). Detecting ^13^C labelled ethanol, which can be present in variable amounts in the background room air, provides evidence that the ethanol was actively produced by microbial metabolism from ^13^C glucose. The amount of variation explained by subject as assessed by Permutational Multivariate Analysis of Variance (PERMANOVA) and was also different for the two different volatile datasets (Table 1, Figure S2). For the ^13^C labeled cultured sputum, 51% of the variation was explained by the subject, while 33% of the variation was explained by subject from the volatiles in the uncultured sputum samples (Table 1). The microbiome community composition as determined by 16S rRNA amplicon sequencing from the seven subjects were unique to each subject (Fig S3), and the individual signatures were also reflected in both the cultured and uncultured sputum volatile molecules detected.

**Table 1.** Permutated multivariate analysis of variance (PERMANOVA) of sputum samples. The PERMANOVA was generated using the “adonis” function from the “vegan” package in R.

**Figure 4.**
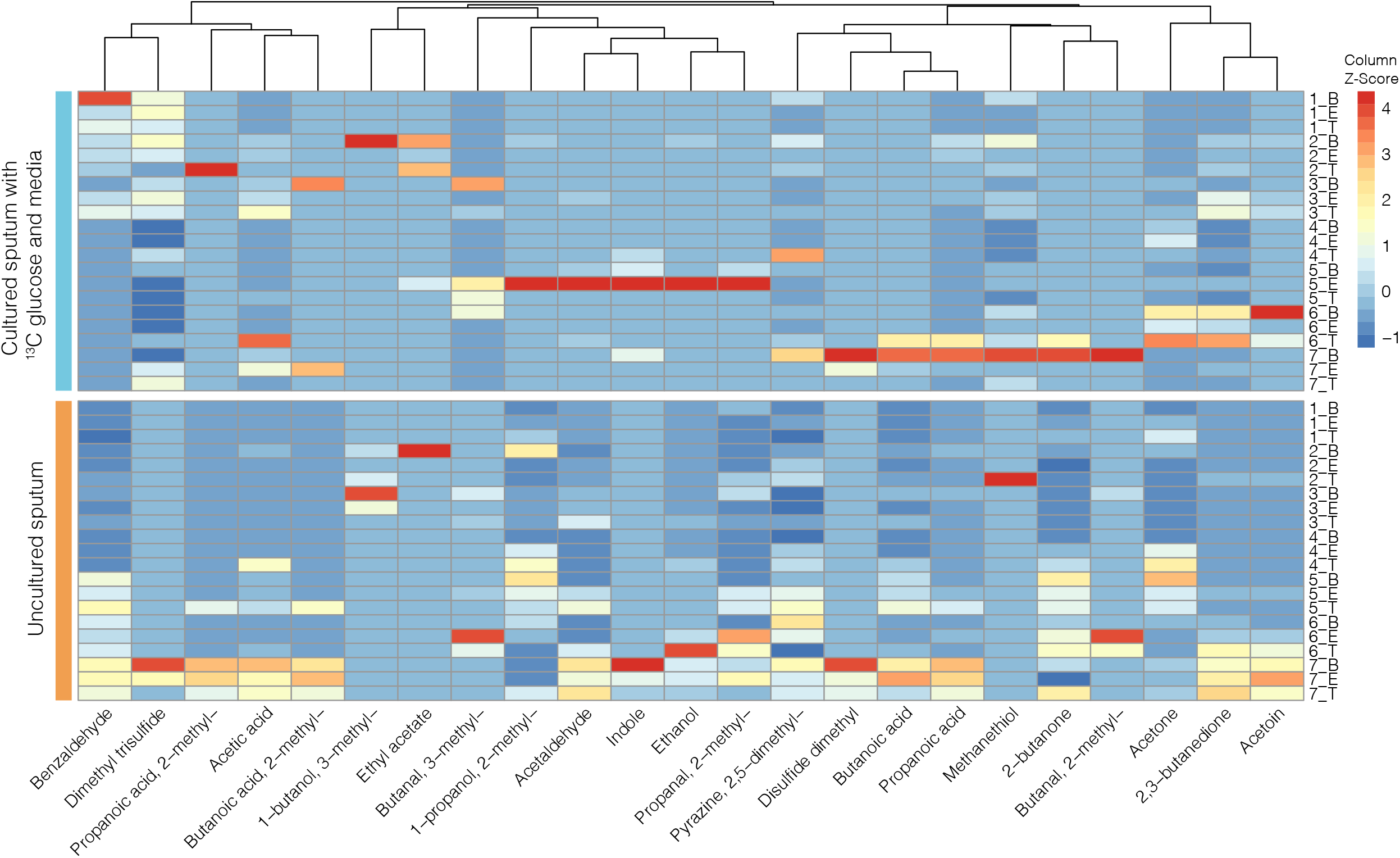
Heatmap of ^13^C labeled volatiles from cultured sputum and volatile molecules detected from uncultured sputum. The labeled volatiles come from the stable isotope probing experiments where ^13^C glucose and BHI were added to sputum during the extraction step to cultivate microbial growth and capture active volatile production. The unlabeled volatile molecules were detected directly from sputum samples. The heatmap intensities are Z-scores as described in Figure 2. However, the Z-scores were calculated separately for the cultured and uncultured sputum experiments. The dendrogram was generated as described in Figure 2.

In cultured sputum, we detected 23 volatiles that were fully labeled with ^13^carbon. The isotope enriched (active) volatiles detected from the sputum samples were different for each subject. The volatiles with isotope enrichment detected in the sputum samples from all seven subjects were 2,3-butanedione; acetic acid; acetone; dimethyl trisulfide; disulfide, dimethyl; and pyrazine, 2,5-dimethyl (Figure 5). Although those volatiles were detected in all subjects, the isotope enrichment for each subject varied. Samples from subject 7 had higher isotope enrichment of disulfide dimethyl compared to the other six subjects (Figure 5B). Acetone was higher in subjects 4 and 6 (Figure 5). Some volatiles were enriched with ^13^C only in certain subjects. For example, 1-butanol, 3-methyl and propanoic acid, 2-methyl were only enriched in a subset of samples from subject 2 (Figure 5). In addition to the isotope enriched volatiles, there were volatiles also detected as unlabeled from the same cultured sputum (Figure S4). Volatiles 2-piperidinone; benzaldehyde, 4-methyl; benzothiazole; butanoic acid, 3-methyl; hexanal; hexane; isopropyl alcohol; phenol; propanoic acid, 2-methyl; and pyrrolo 1,2-apyrazine-1,4-dione, hexahydro were detected in the sputum samples, but were not isotope enriched (Figure S4).

**Figure 5.**
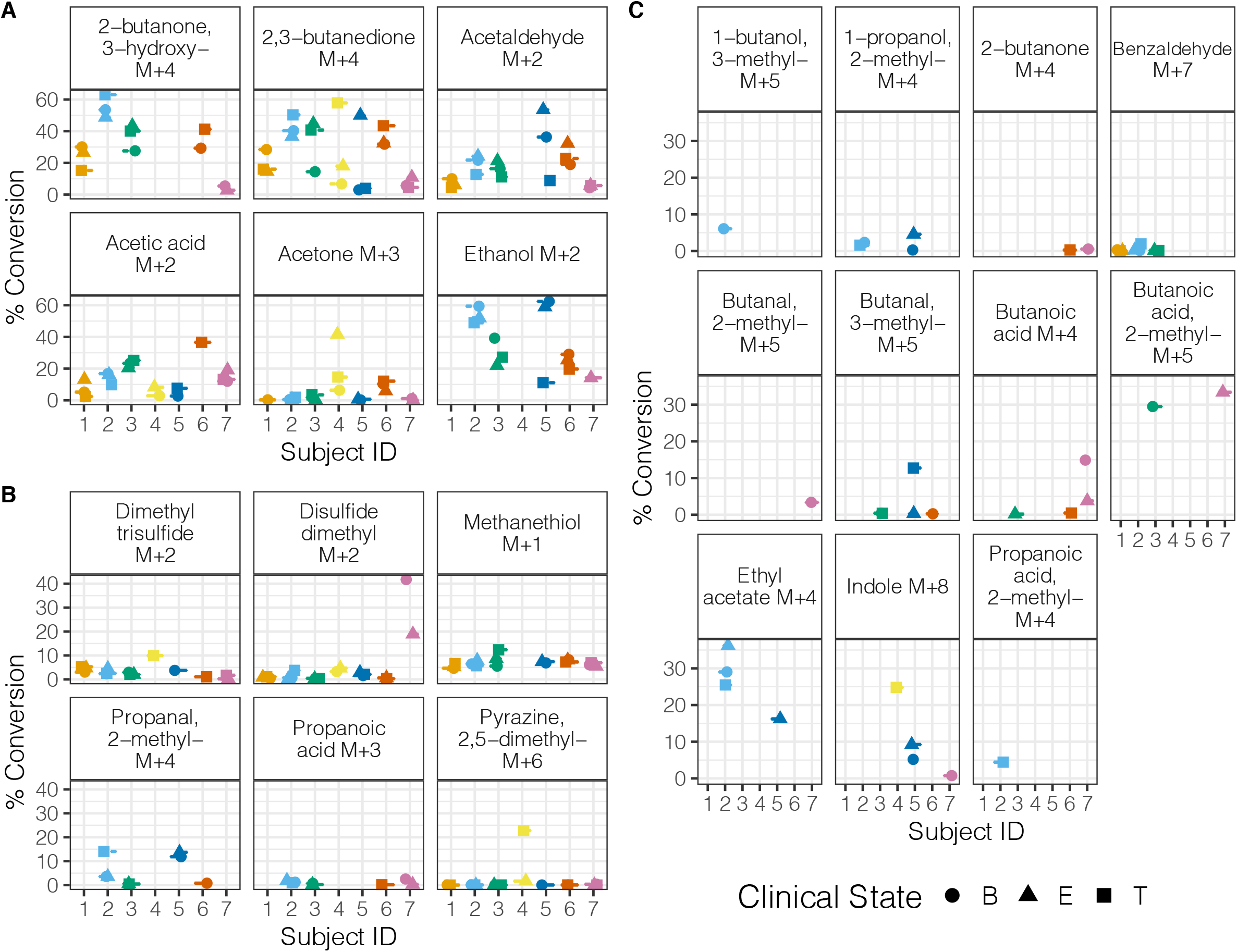
Percent conversion of ^13^C into volatile molecule mass in sputum samples from seven subjects with cystic fibrosis during 18 hours of incubation during extraction. The % conversion was calculated as described in Figure 3. Volatiles not detected in samples are indicated by samples where there are no data. N = 1-3. (A) The ^13^C labeled volatiles detected at a higher percent conversion in the majority of sputum samples. (B) The ^13^C labeled volatiles detected at a lower percent conversion in the majority of sputum samples. (C). The ^13^C labeled volatiles detected at a lower percent conversion in a minority of the sputum samples.

## DISCUSSION

To identify volatile production in *in vitro* cultures and human associated samples, we performed volatile analysis of mono- and co-cultures of *P. aeruginosa, S. aureus*, and *A. baumanii* and stable isotope probing of different biological samples. In the analysis for the mono- and co-cultures, we were able to detect the volatiles present in the samples by performing a short extraction for 1 hour at 70 °C. The volatile analysis of mono- and co-cultures allowed us to survey the volatiles produced by individual species and during their interactions with other species. There were differences in relative abundances across the different culture types and time points. In the stable isotope probing experiments, the biological samples came from feces and saliva from healthy subjects, sewage, and sputum from subjects with cystic fibrosis. Stable isotope profiling enabled us to identify actively produced volatile molecules by extracting for 18 hours at 37 °C. The long extraction time with a lower temperature enabled growth and metabolism of microbes present in the biological samples. Comparing the ^13^C glucose and D_2_O enrichments showed that there was more extensive isotope enrichment with labeled ^13^C.

When extracting different sample types, there were initial optimizing steps to take prior to starting a full run. First, test different volumes of a sample as a trial run. It is recommended to start with a lower volume or smaller amount of sample first. Do not extract a large volume or too much of the sample because the sample could overwhelm the column and contaminate the HSP. Contamination can be evident when peaks in the chromatogram are saturated and there is carryover when the HSP is re-run on the GC-MS. In the mono- and co-culture experiments, 200 μl of culture was sufficient to detect a variety of volatiles. In the stable isotope probing experiments, depending on the sample type, the volume for each experiment ranged from 500 μl to 1 ml. Second, depending on the compounds of interest and column type, the GC-MS and Entech methods will need to be adjusted to optimize volatile detection. We determined our methods appropriate for our analytes of interest and column type.

After optimizing the method, the critical steps in the protocol pertained to the steps prior to and following extraction. During sample preparation, samples were placed on ice so that the volatiles present in the sample did not escape. It was also important to make sure the vacuum seal was tight and lid was securely closed. Otherwise, there would be an inefficient extraction and decreased detection of the volatiles from the sample. A reason there would be a leak in the seal is the O-ring around the lid liner or the pen itself. To make sure that the vial was under vacuum each time, a gauge was used to prior to extraction to make sure the O-rings were still functional. In addition, after the extraction, samples were placed on the ice block so that water was drawn out of the headspace for a determined time period. Water in the column could lead to changes in retention times of the volatile signals.

There are both limitations and advantages to these methods with respect to existing or alternative methods. Evaluating the mono- and co-culture experiments, there were volatile signatures detected that were specific to a particular microbe or co-culture. There were also changes in volatile abundances across time. As for the stable isotope probing experiments in the different types of biological samples, deuterium labeling did not result in as many isotope enriched volatiles as ^13^C glucose. The metabolisms required to produce isotope enriched volatiles with deuterium may be more limited. In addition, media was added to the biological samples to enhance microbial growth, which may lead to changes in the microbial community composition. Culture conditions of stable isotope probing experiments may be selecting for the favored growth of certain microbes in a biological sample. This was opposed to the short extraction for one hour at 70 °C designed to detect the volatiles in the community sample before one or few of the microbes begin taking over the community. This method provided a high-throughput solvent-free and vacuum extraction that led to more sensitive detection of low volume samples. The volumes used as input for these experiments ranged from 200 μl cultures to 1 mL of cultured biological samples. In other cases (data not shown), we have extracted biological samples (e.g, sputum and fecal samples) as low as 15 μl or 10 μg.

For future applications of the method, a wide range of sample types could be analyzed with small or limited volumes. Dozens of samples could be extracted at the same time and the run time would depend on the capabilities of the specific GC-MS instrument. In the case of stable isotope probing, when coupled with metagenomic sequencing, there is the possibility of identifying the microbes responsible for the production of the volatile molecules. The metagenome of the biological sample could be sequenced prior to and after extraction with stable isotope probing to identify changes in microbial community composition. The metagenomic sequencing will allow identification of genes responsible for the production of the isotope enriched volatile molecules. Here, we highlighted a few examples of the sample types and approaches that could be used as input for this method, which has already been established in different industries. Because volatile molecules are important diagnostic indicators, the use of this method could be expanded to biological laboratories and clinical healthcare settings. Overall, there are a great number of possibilities that can result from the use of vacuum assisted sorbent extraction.

## ACKNOWLEDGMENTS

We thank Heather Maughan and Linda M. Kalikin for careful editing of this manuscript. This work was supported by NIH NHLBI (grant 5R01HL136647-04).

## FIGURE AND TABLE LEGENDS

**Figure S1**. The relative abundances of labeled (M+N(max)) and unlabeled (M+0) volatiles across fecal, saliva, and sewage samples.

**Figure S2**. Non-metric multidimensional scaling of cultured sputum with stable isotope probing and uncultured sputum. (A) The NMDS of cultured sputum with ^13^C glucose and media was generated with k = 3 dimensions. The stress value was 0.07. (B) The NMDS of uncultured sputum was generated with k = 3 dimensions. The stress value was 0.13.

**Figure S3**. Microbial community composition of sputum samples from subjects with cystic fibrosis, as assessed by 16S rRNA amplicon sequencing as part of a larger study, further information about approach found in Carmody et al 2020^19^. from subjects with cystic fibrosis. B = baseline, E = exacerbation, T = treatment. Each stacked bar is a different timepoint.

**Figure S4**. The relative abundances of labeled (M+N(max)) and unlabeled (M+0) volatiles across sputum samples from seven subjects with cystic fibrosis.

