## Supplementary figures and images for "Capturing actively produced microbial volatile organic compounds from human associated samples with vacuum assisted sorbent extraction"

### Supplemental Figure 1

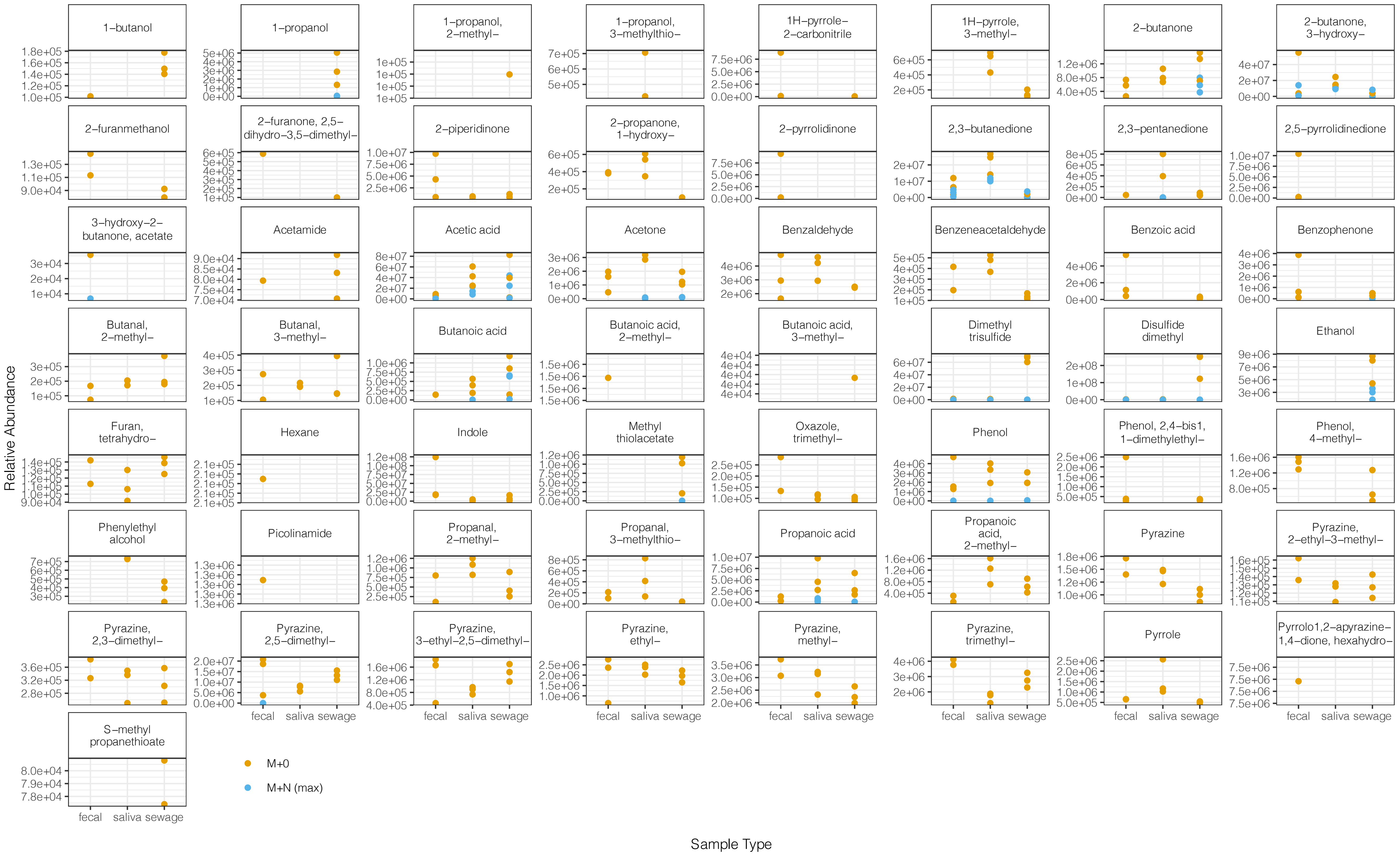

### Supplemental Figure 2

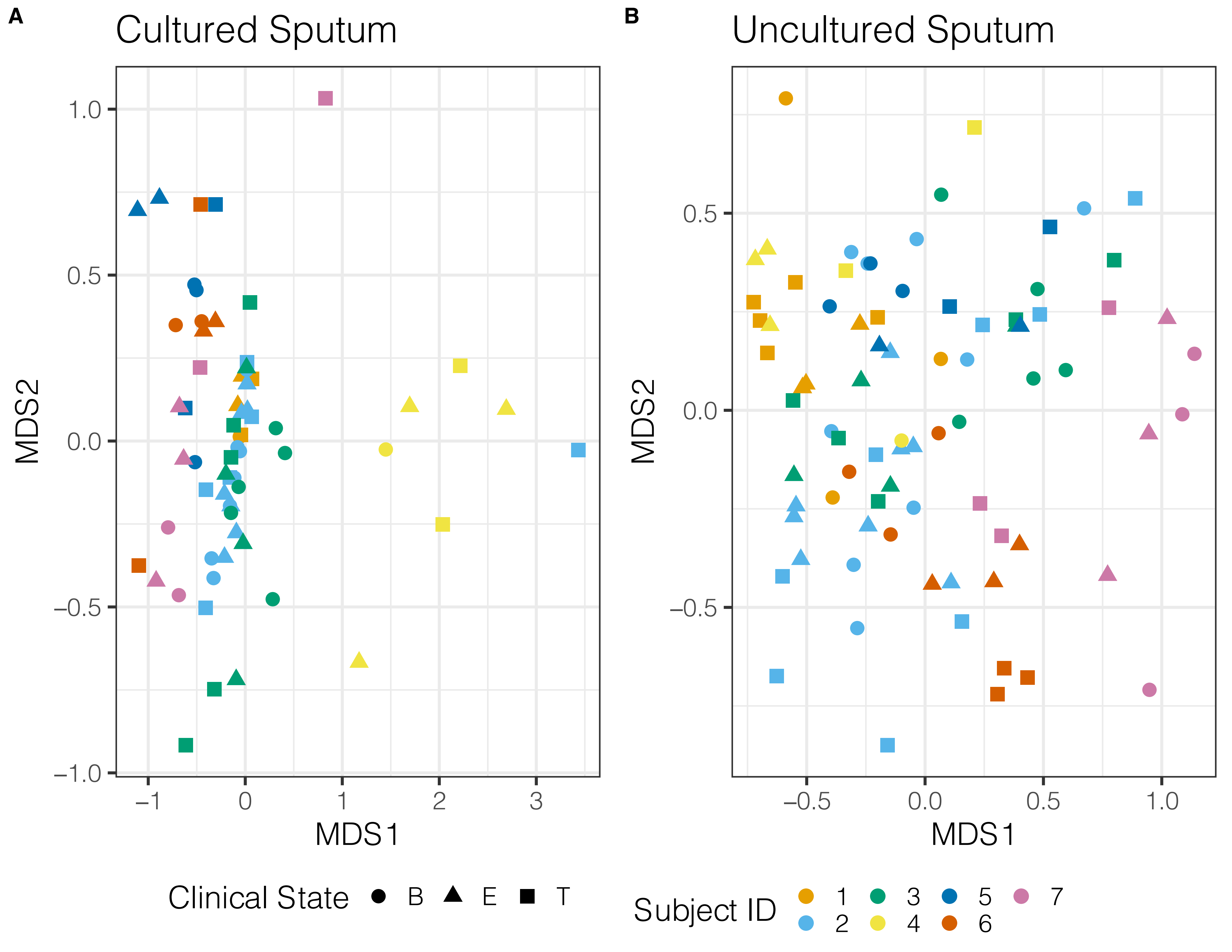

### Supplemental Figure 3

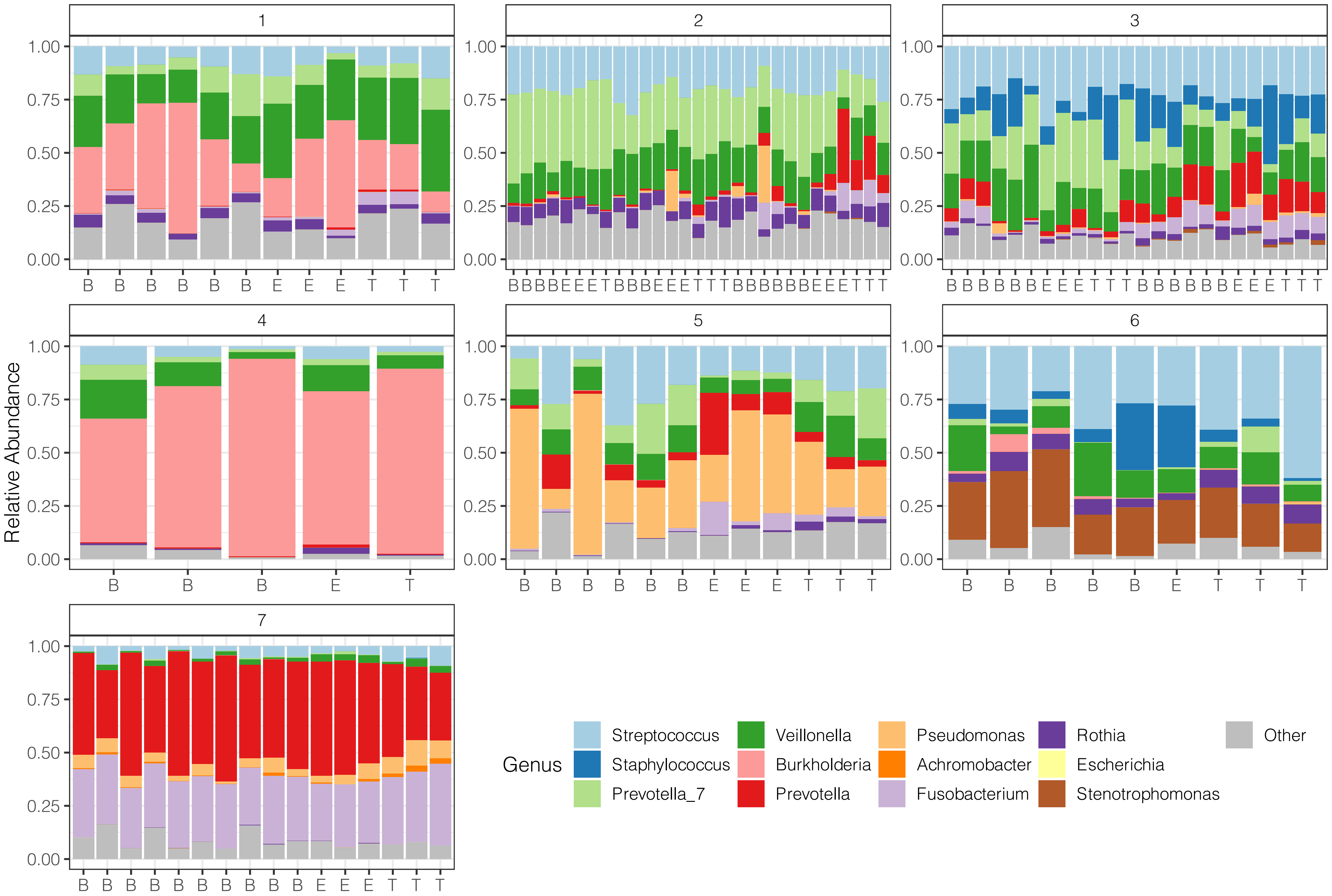

### Supplemental Figure 4

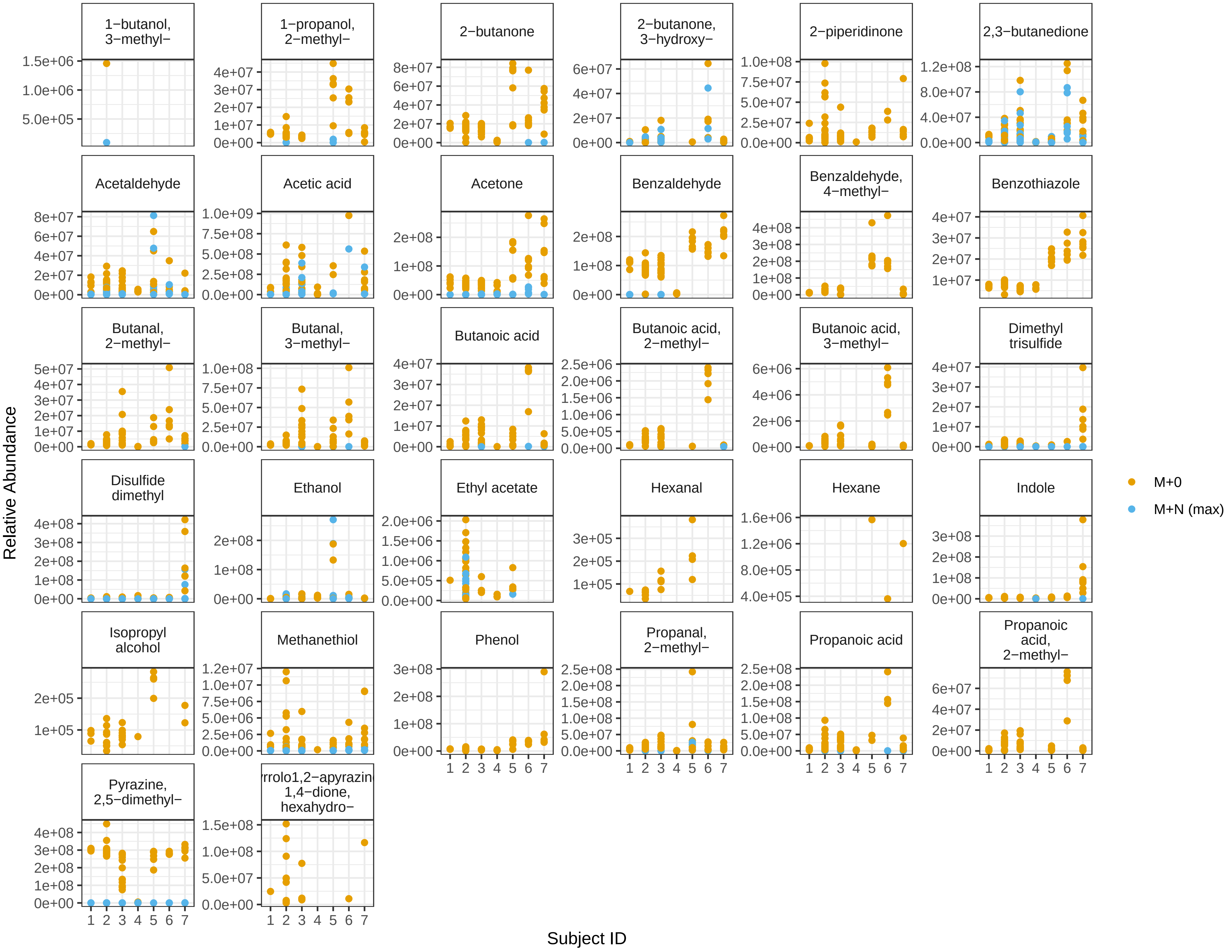
