## Appendix for "Capturing actively produced microbial volatile organic compounds from human associated samples with vacuum assisted sorbent extraction"

**PROCEDURE**

1. **Headspace sorbent pen (HSP) and sample run considerations**
   1.1 The HSP with Tenax TA was selected to capture a broad range of volatiles in consideration with the DB-624 column installed in the GC-MS.
   1.2 Extract media or sample blanks with the same conditions for the same period as sample extraction.
   1.3 Run a blank pen on the GC-MS before running extracted samples using the same method. Blanks should also be run between sample types (e.g. three replicates of bacteria mono-culture, blank, three replicates of bacteria co-culture, blank, etc.)
   1.4 Limit use of fragrant personal care items or consumption of smelly foods prior to sample extraction and analysis. Ideally, samples will be prepared in a biosafety hood that has not been cleansed by alcohol or other volatile cleaners for at least half an hour. Turn on airflow in biosafety hood for half an hour prior to sample preparation.
   1.5 Keep samples on ice to limit volatile release during sample preparation.

**2. Mono- and co-culture preparation**

2.1 Inoculate cultures of *A. baumannii, S. aureus,* and *P. aeruginosa* in Todd Hewitt growth media. Incubate overnight at 37 ℃ with 200 rpm agitation.

2.2 Dilute each culture to optical density 0.05 at 500 nm.

2.3 Mix co-cultures in equal parts and pipet 200 μl of control media, mono-, or co-culture into each well of a 96-well plate and place in 37 ℃ incubator for 24 hours. Prepare a second plate for a 48-hour incubation.

2.4 At the end of the incubation period, prepare samples for extraction in Step 4. Note: at this point, samples can be stored at -80 ℃ to extract later if needed. Pipette liquid cultures into Eppendorf tubes and store at -80 ℃.

**3. Stable isotope probing in biological samples preparation**

**Note:** The feces and saliva samples were donated from anonymous donors with approval from the University of California Irvine Institutional Review Board (HS# 2017-3867). The sewage came from San Diego, CA. The sputum samples were collected from subjects with cystic fibrosis as part of a larger study approved by the University of Michigan Medical School Institutional Review Board (HUM00037056).

3.1.1 To prepare fecal samples, add one mL DI water to 100 mg of feces in a 1.5 mL microcentrifuge tube and vortex for three minutes. Place on ice when not in use.

3.1.2 To 15 μl of fecal and water mixture, add 485 μl Brain Heart Infusion (BHI) medium with 20 mM ^13^C glucose, or BHI with 30% deuterium (D_2_O). The final volume of the sample should be 500 μl. Prepare samples in technical triplicates.

3.2 To prepare sewage samples, add 500 μl of sewage to 500 μl BHI and 20 mM ^13^C glucose or BHI with 30% D_2_O for a total volume of one mL. Prepare samples in triplicate. Place on ice when not in use.

3.3 To prepare saliva samples, add 50 μl of saliva to 500 μl BHI and 20 mM ^13^C glucose or BHI with 30% D_2_O for a total volume of 550 μl. Prepare samples in triplicate. Place on ice when not in use.

3.5.1 To prepare sputum samples to compare the volatiles present in the sample prior to and after culturing, perform a first extraction with 15 μl of sputum. Prepare samples in triplicate. Place on ice when not in use. Proceed to step 4 for sample extraction and extract for 18 hours at 37 ℃ with 200 rpm agitation.

3.5.2 After the completion of the first extraction of the uncultured sputum samples, save the vials with sputum. Add 500 μl BHI with 20 mM ^13^C glucose to the vials with sputum from 3.5.1. Place on ice when not in use.

3.6 Proceed to Step 4 for sample extraction.

**4. Sample Extraction**

4.1 Place empty volatile organic analysis (VOA) vials (20 mL) on the cold plate and place the cold plate on ice.

4.2 Turn on the 5600 sorbent pen extraction unit (SPEU) to warm up to the desired temperature as required for each method. For stable isotope probing experiments at 37 ℃, reaching the setpoint can take up to 15 minutes. For mono- and co-culture experiments at 70 ℃, reaching the setpoint could take up to 45 minutes to an hour.
4.3 Collect clean HSPs that are equal to the number of samples prepared, including HSPs for media or sample controls.
4.4 Label 20 mL VOA vials according to samples, replicates, and pen IDs as needed. It is recommended to use a marker that resists water in case condensation forms on the outside of the vial while on ice.
4.5 Unscrew the white cap on the vial, quickly pipet sample into the vial, and assemble the black cap, lid liner, and HSP. NOTE: Samples should not come into contact with the HSP and sample volume will depend on sample type.
4.6 Place the vial with sample back on the cold plate.
4.7 Repeat steps 4.5 and 4.6 for each sample. These steps are done per sample instead of all at once to prevent sample warming and thus premature volatile release.
4.8 Once all samples have been prepared in the glass vials, turn on the vacuum pump, place the vials under vacuum to 30 mmHg, and remove the vacuum source. The vials do not need to be on the cold tray at this point.
4.9 Double check the pressure after placing all samples under the vacuum using the gauge pressure. If a vial has a leak, ensure that the cap is screwed on tightly and that the white O-rings of the HSP and lid liners are properly in place. A compromised seal can result in a decreased volatile detection compared to volatile detection from a vial under vacuum.
4.10 Place vials in the 5600 SPEU for optimized time and temperature with agitation at 200 rpm. Cultures were extracted for one hour at 70 ℃, and stable isotope probing experiments with fecal, sewage, saliva, and sputum samples were extracted for 18 hours at 37 ℃.
4.11 Place cold plate at -80 ℃ for use after the extraction period is complete.
4.12 When extraction is complete, place samples on cold tray for 15 minutes to draw out water vapor from the HSP and headspace.
4.13 Transfer the pens to their sleeves. Note: The experiment can be paused here for up to ~1 week at room temperature before losing volatiles on the HSPs.

**5. Analyze samples on the GC-MS.**

5.1 Agilent GC-MS (7890A GC and 5975C inert XL MSD with Triple-Axis Detector) with a DB-624 column settings are set as follows: 35 ℃ with a 5 minute hold, 10 ℃/min ramp until 170 ℃, and a 15 ℃/min ramp until 230 ℃ with a total runtime of 38 minutes.

5.2 Set the Entech method with preheat time for 2 minutes and temperature at 70 ℃, desorption for 2 minutes at 260 ℃, bakeout for 34 minutes at 260 ℃, and post bake for 2 minutes at 70 ℃.

5.3.1 Set up sequence of samples and start the run according to instrumentation.

5.3.2 To set up sequence on the Entech Software, open the program. Next to the instrument dropdown menu, click on “5800” and then click on “Sequence” in the options to the right of the drop-down menu.

5.3.3 There is a sequence table similar to the GC-MS side. The “Sample ID” column is named according to “Current date_vial number”, “Name” is the same name of sample as inputted on the GC-MS side, and “5800 Method” determines the rate of temperature ramp, holding times, etc. (opens a menu to select a method).

5.3.4 The “Tray” and “Position” columns determine where the autosampler will go to pick up the VASE HSPs. To the immediate left are two trays with 30 spots each, laid out as six columns with five spots each. The spot furthest and leftmost on each tray is spot 1, while the closest and rightmost spot is spot 30. These trays are “HSP A or B”, where HSP B is the tray closer to the autosampler (innermost tray). Directly behind HSP B is HSP Blank. Place extracted samples into the trays and select the spot on the sequence accordingly.

5.3.5 Save sequence table, click on “Run” on the left-hand side, then “Start with blank in desorber” if the blank HSP is in the desorber (denoted by a HSP marked by yellow label).

5.4 HSPs will be handled by the Sample Preparation Rail (SPR) for each sample in the sequence in the run. The SPR will warm up, then a message will appear at the top of the screen to confirm if the blank is in the desorber. Click on “Skip” to confirm that the pen is there. The autosampler will now run all samples automatically, and the sequence on the GC-MS side will automatically record the data in separate files.

**6. Data analysis**

6.1.1 Quality filter data on GC-MS ChemStation software.

6.1.2 Review each peak on the chromatogram and annotate peaks tentatively identified by the National Institute of Standards & Technology (NIST) library (or with an available library).

6.1.3 To add a peak to the process method, click -> “Calibrate” -> “Edit Compound” -> “Name” -> insert compound under “External Standard Compound”. Add the name of the compound, retention time, Quant Signal “Target Ion”. Add the three largest peaks. To save, click on “ok” -> “Method” -> “Save.”

6.1.4 Once the process method is set up, go to “Quantitate” -> “Calculate.” Next, go to “View” -> “QEdit Quant Result.”

6.1.5 Go through each compound and make sure that the peaks align and are above background noise.

6.1.6 Once QEditing is finished, click on “Exit” -> “Yes” to save the QEdits and return to the main chromatogram. Export the area integrations by opening the file on the left-hand side. Go to “Quantitate” -> “Generate Report.”

6.1.7 To export files to use in DExSI, go to “File” -> “Export Data to AIA format” -> “Create New Directory” and select a location for the file or “Use Existing Directory.”

6.1.8 Another new window will open up to select files for export. Move the files to the right side of the window, then click on “Process.” This conversion may take a few seconds to a few minutes depending on the number of files being converted.

6.2 Correct for isotope abundance in DExSI according to instructions for the DExSI software (<https://github.com/DExSI/DExSI>) and perform analysis with a favorite software or program (e.g., R).
